# Lipopolysaccharides nanodiscs, a biomimetic platform to study bacterial surface

**DOI:** 10.1101/2025.02.05.636646

**Authors:** Abbas Massilia, Micciulla Samantha, Teulon Jean-Marie, Maalej Meriem, Trembley Macha, Marchetti Roberta, Molinaro Antonio, Thépaut Michel, Fieschi Franck, Pellequer Jean-Luc, Laguri Cédric

**Affiliations:** Univ. Grenoble Alpes, CNRS, CEA, Institut de Biologie Structurale, F-38000 Grenoble, France; Univ. Grenoble Alpes, CNRS, LIPhy UMR 5588, Grenoble, France; Department of Chemical Sciences, University of Napoli Federico II; Institut Universitaire de France (IUF), Paris, France

**Keywords:** LPS surfaces, polymyxin, macrophage galactose lectin, QCM-D, styrene-maleic acid copolymers

## Abstract

Lipopolysaccharides (LPS) are essential components of the outer membranes of Gram-negative bacteria, playing a crucial role in antimicrobial resistance, virulence, and the host’s immune response. Self-assembled particles displaying LPS are essential for biophysical studies addressing the behavior of bacterial surfaces under specific biomimetic conditions. Styrene-maleic acid (SMA) copolymers were employed to form LPS nanodiscs, either from extracted LPS or directly from purified outer membranes. These nanodiscs, derived from pathogenic O157:H7 or laboratory *E. coli* strains, are well-defined in size and yield high-resolution nuclear magnetic resonance (NMR) spectra. They have been successfully used to investigate molecular recognition by a human C-type Lectin Receptor (CLR) of the immune system and interaction with polymyxin antibiotics using various biophysical methods. This study highlights the significance of LPS nanodiscs as bacterial surface mimetics and opens promising avenues for further research into LPS structure and interactions. The ability to generate well-defined LPS nanodiscs offers a powerful tool for studying the molecular mechanisms underlying bacterial pathogenesis and immune response.

## INTRODUCTION

The Outer Membrane (OM) of Gram-negative bacteria is essentially built up of LipoPolySaccharides (LPSs), a key glycoconjugate component of its outer leaflet ^1^ (Figure 1). LPSs display chemical and structural variability depending on the bacterial species and the growth conditions. Their structure is constituted of three main parts: the lipid A, formed by N- and O-acylated di-glucosamine; the core oligosaccharide (OS), a region composed of 10-15 sugars; the O-antigen (O-Ag), a polysaccharide that can span up to 20-30 nm. Lipid A together with core OS are the most structurally conserved parts of LPS, while the O-Ag is highly variable, with more than 180 different O-Ag structures referenced for *Escherichia coli* ^2^, for instance. Gram-negative bacteria can produce incomplete LPSs lacking the O-antigen domain, therefore only composed of OS and lipid A moieties. These are referred to as LipoOligoSaccharides (LOS). As schematized in Figure 1, the LPS structures present at the bacterial surface form a heterogeneous mixture of LOS and LPS with variable O-antigen lengths, creating a complex glycan landscape.

**Figure 1.**
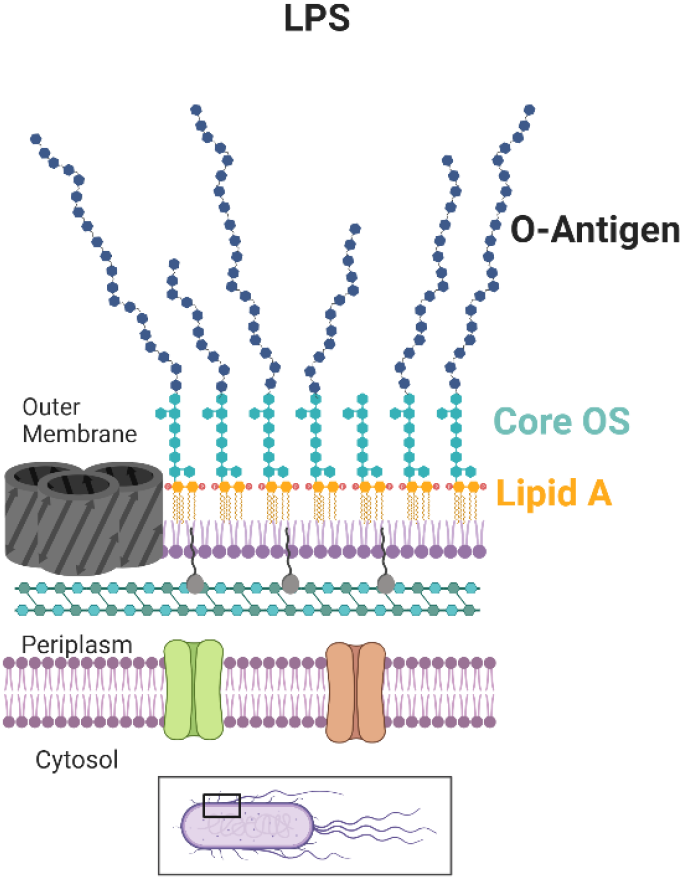
Overview of Gram-negative bacteria cell envelope showing the LPS embedded into the outer leaflet of the OM. Created in https://Biorender.com

From the biological perspective, LPSs have a dual role: 1) they form an efficient barrier, with their lipidic part forming a highly impermeable surface against entry of small molecules, and the O-Ag assembling into a thick sugar layer and contributing to outer-membrane stiffness^3^; 2) they are endotoxins, as they activate the immune system of plants, animals, and humans^4^. The study of LPSs remains challenging due to their heterogeneous and highly variable structure. LOS and LPS can be extracted from bacteria^5^, but assemble into different macromolecular structures in solution depending on the size of the glycan moieties. Indeed, LOSs assemble as vesicles, while LPSs form tubular micellar structures^6^. Given their amphiphilic nature, LPSs purification and chemical fragmentation of their different moieties is necessary for their characterization^5^. However, fragmented structures lose some of their native properties, making them inadequate model systems of real bacterial surfaces. Lipid nanoparticles, also termed nanodiscs (ND), have revolutionized the study of membrane proteins using mostly membrane scaffold proteins^7^. They also hold potential applications for the reconstitution of LPS surfaces at solid and air/liquid interfaces^7^. To generate lipid nanodiscs, styrene maleic acid copolymers (SMA) have been shown to spontaneously insert into native membranes and form lipid-nanodiscs without the prior use of detergents^8^. Here, we have successfully used the SMA system to form NDs from purified LPSs, and also directly from bacterial outer membranes (OM) of both pathogenic and model *E. coli* strains. We have demonstrated how ND-LPS can serve as biomimetic platforms, such as flat bilayers, to study LPS interactions under a variety of biologically relevant conditions, such as the interaction with a human C-Type Lectin Receptor involved in innate immunity and antibiotics that target LPS.

## RESULTS and discussion

### Formation and characterisation of LPS nanodiscs from pathogenic O157:H7 *E. coli*

LPS from pathogenic *E. coli* O157:H7 was first investigated to validate the formation of LPS NDs with SMA polymers on a biologically relevant system. This enterohemorrhagic strain is responsible for regular foodborne outbreaks and its O-antigen is essential for virulence^9,10^. O157 LPS was first purified from bacteria using the Phenol/Chloroform/light Petroleum method^11^. When extracted from the phenol phase, it assembles into large and heterogeneous membrane structures (Figure 2A). Purified O157 LPS was then incubated with SMA S200, and remaining membranes were pelleted by ultracentrifugation. The supernatant, containing nanodiscs, was subjected to a sucrose gradient to remove free polymers (see Methods section) and the samples are referred to as ND-0157_pur_. Negative-stain transmission electron microscopy, Dynamic Light Scattering (DLS) and atomic force microscopy (AFM) confirmed the successful formation of nanodiscs with a relatively homogeneous size distribution (Figure 2, A-B Figure S1).

**Figure 2.**
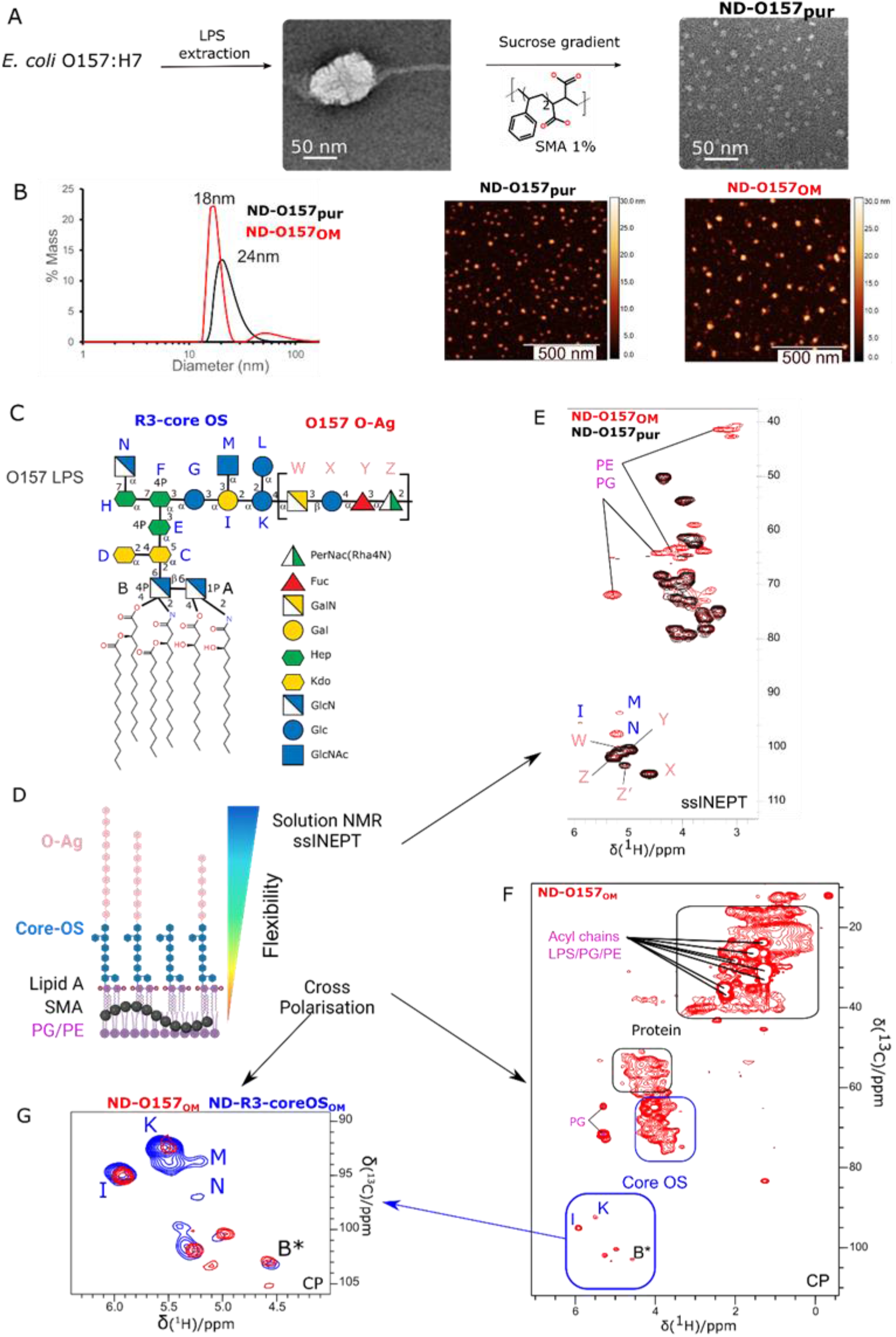
Preparation and characterization of LPS nanodiscs. A) Negative stain electron micrographs of purified O157 LPS from the phenol phase(left) and after SMA solubilization showing ND-O157_pur_ nanodiscs (right). B) Characterization of ND-O157_pur_ and ND-O157_OM_ by DLS (left) and AFM imaging (right). C) Chemical structure of *E. coli* O157 LPS with sugars represented in standard glycan nomenclature snfg^41^. D) Scheme of NMR characterization of ND-O157 depending on the relative flexibility of the different parts of the system, created in https://Biorender.com. E) Overlay of ssINEPT ^1^H-^13^C spectra of ND-O157_pur_ and ND-O157_OM_. F) Cross polarisation of ND-O157_OM_ with the different component of OM components highlighted. G) Comparison of core OS signals in ND-R3_OM_ and ND-O157_OM_. * assessed from _18,19_

Using a different approach, LPS NDs were formed by directly incubating purified outer membranes of *E. coli* O157:H7 with SMA polymers, the polymer ability to extract lipids directly from the native membranes ^12^. In this approach, bacterial disruption was followed by the separation of outer and inner membranes *via* a sucrose gradient^13^. Isolated outer membranes were then treated with SMA polymer, and the resulting NDs were isolated as described above for purified LPS. These samples are referred to as ND-0157_OM_. DLS measurements showed the formation of a majority of NDs with a mean diameter of 18 nm with lower polydispersity compared to ND-O157_pur_ (Figure 2B). Given the low contrast of NDs in negative-staining Electron Microscopy, the samples were imaged by AFM. NDs were adsorbed onto a mica surface coated with Ni^2+^ ions and imaged in air under ambient conditions. AFM images showed well-defined nanodiscs for both ND-O157_OM_ and ND-O157_pur_ (Figure 2B, Figure S1). The formation of LPS NDs with SMA is thus effective and allows the formation of NDs of about 20 nm diameter.

### NMR investigation of ND-O157

NMR spectroscopy is a powerful technique to study structure and interactions of LPS-containing systems. It is non-destructive and allows probing molecular structures at the atomic scale. For this purpose, ^13^C is necessary for a gain in sensitivity and the possibility to acquire NMR correlation experiments through ^13^C nuclei. NDs were thus formed with LPS from bacteria grown in minimal medium with ^13^C glucose as sole carbon source^14^. ND-O_157pur_ were analyzed first by solution NMR. Because of high thermal stability of SMA nanodiscs^8^, experiments could be recorded at 50°C for favorable molecular tumbling. The resulting NMR spectra show with high sensitivity the signals of the O-antigen tetrasaccharide repeat of O157 (Figure 2C) which is highly flexible (Figure S2). In contrast, moieties closer to the membrane (Lipid A and core OS) were not observable due to the slow molecular tumbling of nanodiscs in solution. Magic Angle Spinning (MAS) NMR, on the other hand, enables to average strong couplings that limit solution NMR of large macromolecules^6,15. 13^C-labeled NDs were sedimented inside 1.3 mm rotors and studied at 55 kHz rotation with ^1^H detected experiments through two types of correlation transfer, cross polarization (CP) or insensitive nuclei enhanced by polarization transfer (INEPT). These experiments are overall sensitive to rather rigid or more mobile parts of the system, respectively (Figure 2D). ND-O157_pur_ ^13^ C-^1^H INEPT is identical to the spectrum in solution (Figure S2) and shows the O-antigen as well as the CH_2_s and CH_3_ of the extremity of the Lipid A acyl chains, that are still very dynamic inside the membrane^6^. The O-antigen ^1^H and ^13^C resonances could be assigned under MAS by ^1^H detected and ^13^C detected experiments (Figure S3).

ND-O157_OM_ ^13^C-^1^H INEPT spectrum shows perfect superimposition of the O-antigen signals with ND-O157^pur^ (Figure 2E). As expected, ND-O157_OM_ shows additional signals corresponding to OM components. Phospholipids head groups from Phosphatidyl-Ethanolamine (PE) and Phosphatidyl Glycerol (PG) were also observed, as well as lipid modifications such as unsaturation of fatty acids^6^. (Figure 2E, Figure S4). ^1^H-^13^C CP spectra, on the other hand, tend to select rigid parts. O157 O-antigen is filtered out and the CP ^1^H-^13^C spectrum displays signals characteristic of proteins, that were coextracted with LPS (Figure 2F, Figure S4). While the spectrum of outer-membrane NDs is complex due to its heterogeneous composition, the sugar anomeric resonances (H_1_-C_1_) were well resolved in the 90-105 ppm region in ^13^C (Figure 2F, G). NMR assignment of these sugar resonances using CP-based experiments was unsuccessful due to low sensitivity. However, they presumably arise from the core OS and Lipid A sugars. O157 core oligosaccharide is of R3 type^16^ (Figure 2C). Therefore, NDs from the outer membrane of F653 *E. coli* strain, that produces LOS containing the R3 type core oligosaccharide, were produced. ^13^C-^1^H INEPT spectrum of ND-R3_OM_ is superimposable to ND-O157_OM_ spectra, except for the absence of O-antigen signals (Figure S4). The anomeric region of ND-O157_OM_ and ND-R3_OM_ in CP-based experiments (Figure 2G) largely overlap, confirming the signals arise from the core OS. Some of these core oligosaccharides anomeric groups could be assigned (I, M, N, K) by solution NMR of O157 LPS solubilized in detergent micelles (see methods section) (Figure 2G, Figure S5). Their chemical shifts are very similar to the assignments of R3 core OS from the litterature^17^. Assignment of Lipid A glucosamine (B) remains tentative but corresponds to a highly substituted glucosamine with chemical shifts similar to previous reports ^18,19^. Overall, NMR analysis of NDs consistently shows high resolution spectra either in solution or solid-state NMR depending on the relative flexibility of the different molecular components.

### ND-LPS constitute a relevant platform to explore interactions with a lectin from the immune system

We next investigated the potential use of ND-LPS to reconstitute bacterial surfaces for recognition studies. As a proof of concept, a simple system was chosen, that contains only LOS (without O-antigen). The R1 core oligosaccharide is extensively used to study interactions with proteins^20–23^ and is the major core type found in pathogenic *E. coli* ^16^. Therefore ND-R1_pur_ and ND-R1_OM_ were produced following the same approach as for O157 LPS (see methods section and Figure S6). The human Macrophage Galactose-type Lectin (MGL) was selected as a test protein. This trimeric C-type lectin receptor, expressed in dendritic cells, binds R1 core OS on *E. coli* surface with high affinity thanks to two LPS binding sites per MGL monomer ^23^. To assess this interaction, ND-R1_pur_, composed only of LOS molecules, were first functionalized with a biotin through their maleic acid moieties, as previously described ^24^, and immobilized on streptavidin-coated Biolayer Interferometry (BLI) sensors (see methods section). Injections of increasing concentrations of MGL show a dose-dependent interaction and a very slow dissociation. Association and dissociation kinetics could be fitted to an apparent affinity constant of 4 nM (Figure 3A, Figure S7), consistent with the observation of stable binding of MGL to *E. coli* surface ^23^. As a control, MGL was biotinylated and immobilized on BLI biosensors. Purified R1 LOS, that assemble spontaneously as vesicles of various sizes in solution, were flowed over immobilized MGL (Figure S8). Similarly, dissociation was very slow and an apparent *K*_*d*_ of 129 nM could be determined for MGL-biot:R1 vesicles affinity, much higher than the 3 nM estimated for ND-R1pur-biot:MGL. It can be hypothesized that only a small proportion of LOS are available for binding on the spherical vesicles. This, in turn, would highly increase the apparent *K*_*d*_. Biotinylated ND-R1_OM_ were also immobilized on BLI and tested for MGL interaction. A *K*_*d*_ of 3 nM could be estimated, demonstrating that ND-R1_OM_ can be also used to assess protein binding to LPS.

**Figure 3.**
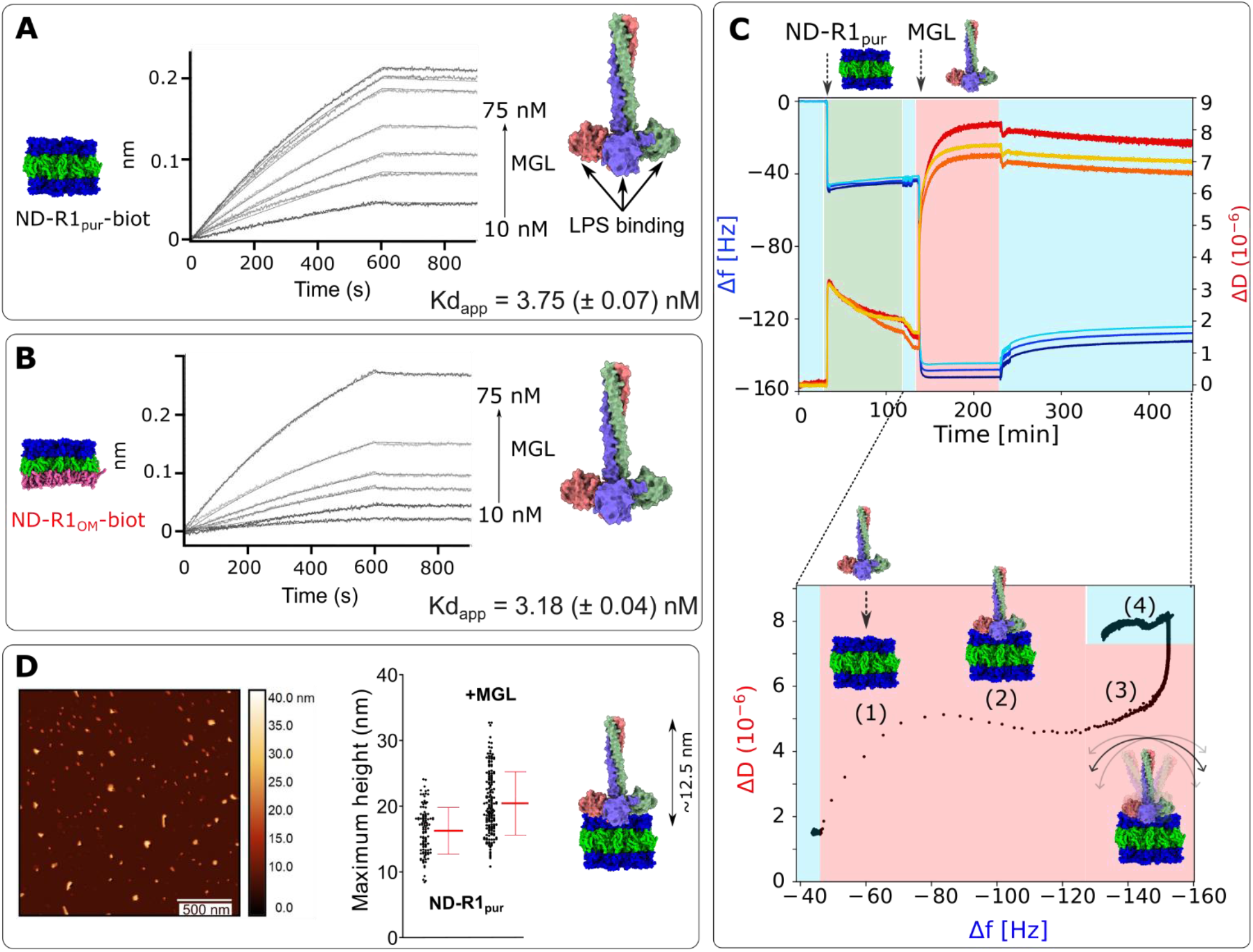
ND-R1_pur_ -MGL Interaction. Biolayer interferometry interaction of immobilized ND-R1_pur_-Biot (A) and ND-R1_OM_-Biot (B) with increasing concentrations of MGL. MGL trimer model^23^ as well as a model of pure LOS or asymmetric LOS bilayers are shown with Lipid A(green), core OS (blue) and phospholipids(pink). C) Frequency (Δf) and dissipation energy (Δf) monitoring for the 5^th^, 7^th^ and 9^th^ overtones during ND-R1_pur_ adsorption and MGL binding with time (left). The rinsing steps (buffer) are marked with a cyan background. ΔD vs Δf plot of MGL adsorption and rinsing phase with proposed schemes of structural behavior at the interface. D) AFM imaging of ND-R1_pur_ upon incubation with MGL (left) with corresponding heights distribution (right). MGL model is shown at the same scale with a symmetrical R1 LPS bilayer.

These results demonstrate that LOS nanodiscs are valuable platforms to assess the kinetics of protein LPS interactions by BLI. However, more accurate structural information on the protein-lipid interaction are beyond the capability of this method. For this purpose we carried out Quartz Crystal Microbalance with dissipation monitoring (QCM-D) experiments to study the properties of ND-R1_pur_ deposited onto a solid substrate and the binding of MGL. QCM-D is a highly sensitive technique to mass variation and viscoelastic properties of thin films adsorbed onto a solid surface. It exploits the piezoelectric properties of quartz to drive a crystal to its resonance frequency and higher overtones applying an alternating voltage across its surface. All the variations of these oscillation frequencies due to the coupling of a mass on the sensor surface produce a change of frequency shift (Δf), which is proportional to the coupled mass (with negative variation for added mass, and vice versa). Moreover, the instrument allows the quantification of the energy dissipated by the bound system (ΔD), whose amplitude changes according to the viscoelastic character of the adsorbed layer. This simultaneous acquisition of the Δf and ΔD at different overtones is crucial to study binding and conformational changes of biomolecular systems, which have in most cases a very pronounced frequency-dependent behavior ^25^. In our experiment (Figure 3C), ND-R1_pur_ were deposited onto a gold-modified quartz crystal, showing a fast adsorption kinetics (green background in Figure 3C) and a very stable binding, with no significant mass loss over time. The measured frequency shift of about -40 Hz is consistent with the deposition of a lipid bilayer. The slightly higher values for both Δf and ΔD (ΔD ∼2 × 10^−6^) compared to canonical phospholipid bilayers (Δf ∼ -30 Hz, ΔD ≤ 1)^26^ can be explained by the higher hydration of sugar moieties of the core OS and the presence of charged polymer units (SMA belt). Under the validity of the Sauerbrey model^27^ and assuming an approximate film density of 1 mg/cm^328^, we obtain a thickness of 7.6 ± 0.2 nm for deposited ND-R1_pur_^29^. This is in good agreement with values reported for supported lipid bilayers and the expected thickness of a pure LOS bilayer (about 8.5 nm). The injection of MGL protein (red background in Figure 3C) generated a significant negative frequency shift (Δf of ∼90 Hz), consistent with a large mass increase upon protein binding. Furthermore, the increase of dissipation energy (ΔD ∼ 7.0 × 10^−6^) is a typical indication of the presence of a highly hydrated and flexible mass. In support to this interpretation, both Δf and ΔD show a constant overtone splitting upon equilibration consistent with an increase in height of the system. Finally, the minor variations measured upon rinsing and long equilibration times showed a strong and stable interaction of MGL with R1 core OSs (Figure 3C, lightblue background), consistent with the stable complexes observed in BLI.

A more accurate interpretation of the adsorption process and possible structural rearrangement at the surface could be extracted from the representation of ΔD versus Δf traces from the MGL adsorption (red) and rinsing (blue) phases (Figure 3C right). Four different phases of MGL behaviour can be observed from ΔD versus Δf traces (1-4 Figure 3C). First, the large separation of the data points at the beginning of MGL injection (phase 1) demonstrates the strong affinity and quick binding of the protein to the LPS nanodiscs, followed by a stabilization of the interfacial mass and a slower filling of the surface (phase 2: Δf decrease, ΔD constant). When the surface saturation is approached, the protein starts a structural rearrangement (phase3), which leads to a significant increase of dissipation energy (visco-elastic behavior) while the adsorbed mass remains constant. This could be the signature of a rotation and oscillation of its elongated, outermost region. At the final rinsing step (phase 4), a minor mass loss was registered, most likely due to loosely bound protein molecules which are rinsed off from the interface, while the viscoelastic character was constant, a clear indication of the MGL stability onto the ND-R1_pur_ surface. It is worth to say that this specific molecular rearrangement is characteristic of MGL only when it binds onto the LPS nanodisc. As a control experiment, we have adsorbed the same concentration of MGL onto a bare gold substrate (Figure S9 A). In this case, while the adsorbed mass is comparable (Δf ∼ -88 Hz), the increment of dissipation is much lower (< 2 × 10^−6^) and no overtone splitting is observed. This means that a rigid mass is bound onto the substrate, with MGL most probably adsorbing in a flat, horizontal conformation, in contrast to the vertically oriented conformation when it binds to the R1 surface. Accordingly, the direct ΔD versus Δf comparison showed a rather simple trend with a fast initial mass increment with low dissipation energy, a constant plateau and a minor dissipation decrease during rinsing.

In conclusion, biolayer interferometry and QCM-D confirm a strong interaction of MGL with LipoOligoSaccharides at the surface of nanodiscs, and, as expected from the localization of MGL sugar binding domains, an orientation of MGL perpendicular to the Nanodiscs surface. Height measurements of ND-R1_pur_ in complex with MGL using AFM further supported this orientation of MGL on LPS surface. The measurements indicated an increase in the mean height of nanodiscs of approximately 4 nm (see Figure 3D and Figure S10). This increase is presumably attributable to MGL-carbohydrate recognition domains (CRDs) that are bound to the surface. Additionally, a substantial rise in the maximum height of the complexes was observed, reaching approximately 8 nm for a maximum size of MGL trimer estimated to be 12.5 nm^23^ (Figure 3D). This finding aligns with the perpendicular binding of MGL on LPS, which exhibits a degree of flexibility, as evidenced by the viscoelasticity determined through QCM-D.

### ND-LPS and antimicrobial peptides interaction

LPS constitute the target of polymyxin antibiotics, positively charged cyclic peptides (Figure 4), which interact with the Lipid A through the phosphate moieties and divalent ions. This family of antimicrobial peptides is used mostly against multi-drug resistant gram-negative bacteria as last resort treatment due to its high toxicity^30^. While the exact mode of action of polymyxins is still controversial, they are able to accumulate at the outer membrane and disrupt LPS assembly, causing cell death^31^. Recent investigations of Polymyxin B (PmB) and its non-lethal variant Polymyxin B nonapeptide (PmBN), that lacks PmB terminal amino acid and acyl chain, (Figure 4) shed light on PmB mechanism. PmB interacts with LPS with high affinity at low nanomolar concentrations until LPS surface is saturated, then binds with lower affinity with fast association and dissociation kinetics^32^. On the other hand PmBN only displays the low affinity component of PmB interaction without accumulation in the LPS surface. This behavior observed by Buchholz and colleagues on *E. coli* cells and outer membrane vesicles by surface plasmon resonance was investigated on ND-R1_pur_ and ND-R1_OM_ by BLI (Figure 3A, B). Consecutive injections of PmB and PmBN were performed on the same biosensors loaded with ND-R1_pur_ or ND-R1_OM_ to observe surface saturation. At low PmB concentrations (33 and 100 nM Figure 4 A,B) no dissociation is observed, consistent with accumulation of PmB at the LPS surface. Higher concentrations injected show low affinity kinetics as the LPS surface becomes saturated with PmB. PmBN, on the other hand, does not show accumulation to the surface and only low affinity kinetics, as previously described^32^. The same kinetics were observed for LPS nanodiscs either purified or extracted from outer membrane.

**Figure 4.**
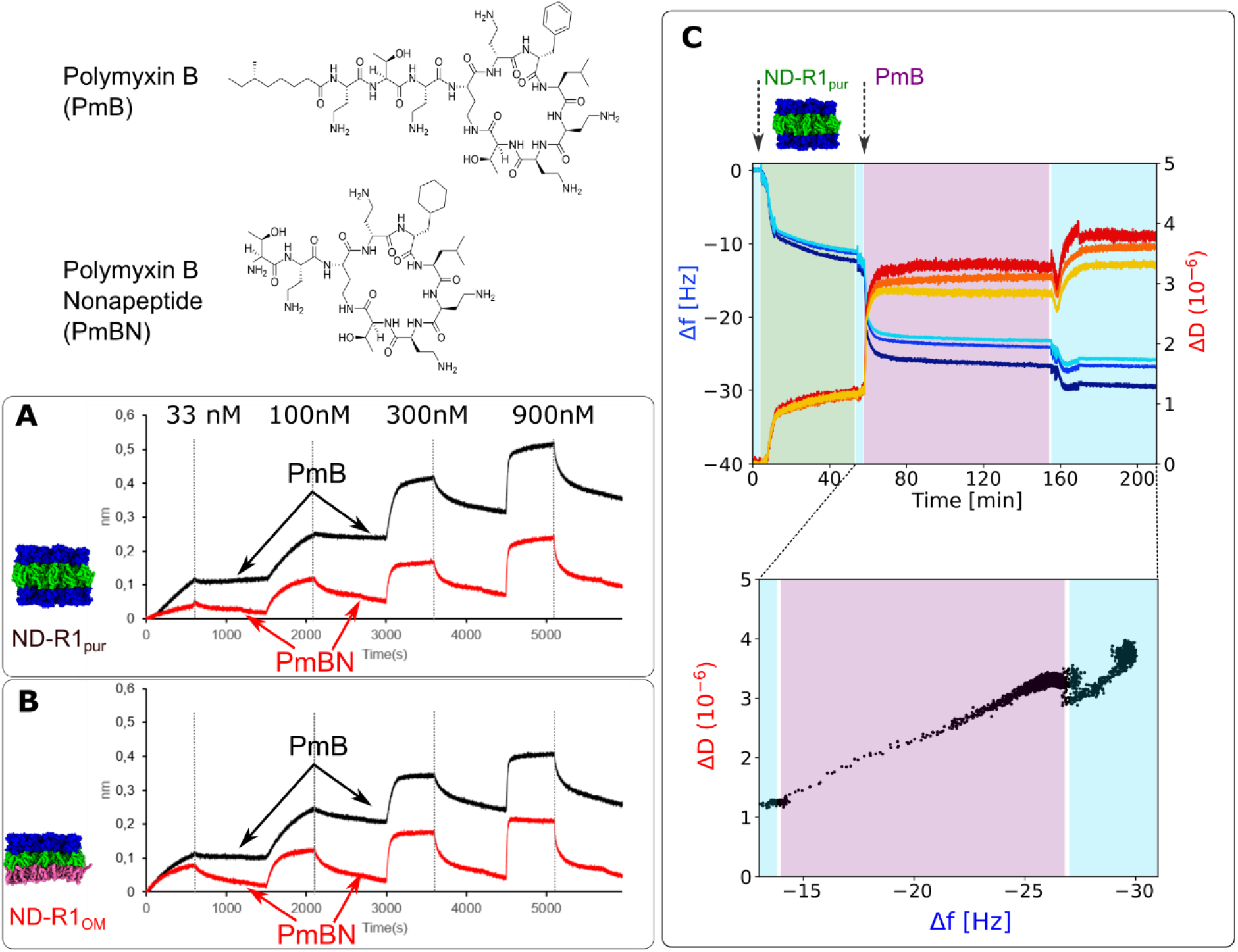
Interaction of ND-R1_pur_ and ND-R1_OM_ with polymyxins. PmB and PmBN are allowed to associate and dissociate at increasing concentrations over immobilized ND-R1_pur_ (A) or ND-R1_OM_ (B). C) Frequency (Δf) and dissipation energy (Δf) monitoring for the 5th, 7th and 9th overtones during ND-R1_pur_ adsorption and PmB binding. The rinsing steps (buffer) are marked with a blue background in the plot.

As for MGL binding, the interaction of PmB, with the LPS layer was assessed by QCM-D. In Fig.4C we report the variation of Δf and ΔD for the adsorption of ND-R1_pur_ (green background), followed by the injection of PmB (purple background). First of all, we can observe in this case a different mechanism of formation for the lipids bilayer, with a slower adsorption kinetics and an overall lower adsorbed mass (Δf ∼ -14 Hz). This could be explained by the different experimental conditions, more precisely the use of a different buffer (see methods). Despite the different binding kinetics and total mass, the QCM-D data confirm the stable nanodiscs binding. The injection of PmB produced an additional mass increment (Δf ∼ -13 Hz), accompanied by dissipation energy increment and overtone splitting. Given the small size of the peptide, the experimental overtone separation could be explained by the creation of defect in the bilayer upon insertion of PmB. This can lead to heterogeneity, and therefore more hydrated domains, or to the reorganization of the polymer belt around the nanodiscs, whose negatively charges might promote some additional interaction with the positively charged peptide. Noteworthy, the total variation of the frequency shift is in this case significantly lower than for MGL, which is reasonable considering the different structure of the two molecules. The rinsing with buffer did not produce a clear mass loss, but apparently an additional increment of the layer hydration. The overall interpretation of LPS-PmB interaction can be further rationalized from the interpretation of the the corresponding ΔD *versus* Δf plot. We can clearly identify the fast adsorption from the frequency and dissipation increment following the PmB injection (purple background), which varies with a comparable rate; this is followed by a stabilization of both signals. The final rinsing step leads to a minor mass and dissipation increment, which is most likely due to water entering some defects in the bilayer. The control experiment for PmB adsorbing onto a bare gold substrate (Fig. S9 B) shows a comparable variation of Δf (∼ -13 Hz), but with a much lower dissipation increment, signature of a flat, rigid layer adsorbed onto the substrate. The final rinsing of the PmB (light blue background) indicates some mass loss with a constant dissipation (Δf ∼ 9 Hz), possibly due to the removal of loosely bound molecules but leaving the surface properties unchanged. ND-LOS are thus able to report interactions with antimicrobial peptides that either interact transiently like PmBN and have poor bacterial killing ability, and PmB that on the other hand are able to insert into the LPS membrane and be bactericidal at low concentration.

### Conclusion

Cell-surface glycoconjugates are ubiquitous and present on almost all cell surfaces, anchored to various macromolecules such as proteins or lipids. In bacteria, lipopolysaccharides (LPS), lipoteichoic acids, and mycobacterial glycolipids are essential components of cellular surfaces. These molecules play both structural and functional roles in resistance to external factors and interactions with hosts. The remarkable chemical diversity of these glycoconjugates and their surface presentation create a heterogeneous three-dimensional mesh, complicating their study *in vitro* under conditions that mimic their natural configuration. The objective of this study was to develop a unified platform for studying lipopolysaccharides of different compositions. LPS from pathogenic bacteria or lipooligosaccharides (LOS) from laboratory strains, purified by solvent extraction, were assembled into nanodiscs upon incubation with styrene-maleic acid (SMA) polymers. Similarly, simple outer-membrane purification followed by incubation with SMA also allowed nanodisc formation. All tested LPS nanodiscs exhibited thermal and long-term stability similar to previously reported lipid-SMA nanodiscs. NMR studies, using ^13^C labelling, provided atomic-scale information on individual glycolipid groups with high resolution through both solution and solid-state NMR in reasonable acquisition times. To test the suitability of these nanodiscs, we chose the simplest and most well-characterized lipooligosaccharides. We investigated the interaction of a human C-type lectin receptor with the LPS, measuring kinetics data, physicochemical behavior using QCM-D, and topography using AFM. These data are consistent with current knowledge of MGL binding to LPS on bacterial surfaces through strong multivalent adhesion, with an orientation of the protein perpendicularly to the membrane^23^. Additionally, we tested the interaction of LPS-assembled nanodiscs with antimicrobial peptides, allowing us to observe non-reversible embedding into the LPS membrane.

The assembly of biologically important glycolipids into a stable platform, accessible to multiple analytical techniques, represents a significant advancement in the study of bacterial surfaces. They allow biophysical study of the LPS as well as interaction with its ligands, not only in bulk but also upon immobilization of solid substrates. Furthermore nanodiscs formed directly from purified outer membranes do not require complex extractions that vary in efficiency depending on the LPS structure, and can be formed from various bacterial strains. Recently SMA NDs from *P. aeruginosa* outer membrane vesicles have also been successfully used to vaccinate mice and monitor phagocytosis^33^. SMA polymers are also easy to functionalize, and immobilize on several supports. More than two dozen polymers capable of forming native nanodiscs have been developed and are commercially available ([SMA Polymers](https://smalp.net/polymers.html)). This allows to easily change the nature of hydrophobic or hydrophilic moieties of the polymer. Polymer change can alter the size of NDs generated but also their susceptibility to divalent ions, the main limitation of the SMA polymers.

One of the most interesting perspectives of our study would be the use of functionalized outer membrane nanodiscs from various pathogenic bacteria to design specific LPS glycan arrays. This would enable to take into account the multivalency of the real bacterial surface to screen for ligands (proteins, antibodies, small molecules). The same NDs could be used to further explore interaction hits with biochemical and biophysical methods.

## MATERIAL AND METHODS

### Bacterial growth

*E. coli* O157:H7 cells (ATCC 43888) and F653 were grown in M9 minimal medium (Na_2_HPO_4_ 5.5 g/l KH_2_PO4 3 g/l, NaCl 0.5 g/l, MgSO_4_ 1 mM, CaCL_2_ 0.1 mM, Thiamine 2mM), enriched with ^15^N-NH_4_Cl (1g/L) U-[^13^C] -glucose (2g/L) at 37°C 200 rpm until mid-exponential phase (OD_260nm_ ∼1). *E. coli* F470 were grown in LB at 37°C, 200 rpm until mid-exponential phase.

### SMA nanodiscs formation and purification

*E. coli F470* (R1) LOS and *E. coli O157:H7* LPS were purified using PCP and phenol/hot water methods, respectively ^14,23^. *E. coli R1* LOS *and O157* LPS at 1.8 mg/mL in buffer A (200 mM NaCl, 50 mM Tris pH 7.4) were incubated under gentle rocking with 1 % SMA S200 (w/v) (Orbiscope) for 30 min at 25°C. Insoluble material was removed by ultracentrifugation at 100 000 g for 1h. NDs were separated from the residual SMA polymer using a sucrose gradient (prepared in buffer A) with three sucrose solutions at 45 % (w/v), 25 %, and 5 %. Sucrose gradients were centrifuged at 100000 g, overnight at 4 °C. Fractions containing NDs were pooled and extensively dialyzed against buffer A, and concentrated using a vivaspin 15, MWCO 100 KDa.

Cells were harvested by centrifugation at 5500 g at 4°C. Cells were lysed using a microfluidizer at 4°C and 15000 psi, debris were removed by centrifugation for 20 min at 10 000 g and total membrane collection by ultracentrifugation at 100000 g for 1h. Inner and outer membranes were separated using a sucrose density gradient: 3 ml 73% (w/v), 4 ml 53% (w/v), 1 ml 20% (w/v). The OM was collected and dialyzed extensively against buffer A. The absence of inner membrane contamination was assessed by the absence of NADH dehydrogenase enzymatic activity^34^. The purified OMs were solubilized using 1% SMA S200 (w/v) for 2 h at 25°C. Insoluble material was removed by ultracentrifugation at 100 000 g for 1h followed by a second sucrose density gradient for residual polymer removal (45% (w/v), 25% (w/v), 5% (w/v)). The fractions containing the nanodiscs were collected, dialyzed against buffer A and concentrated. *E. coli F470* cells were grown in LB medium till mid-exponential phase (2.5 OD_260nm_). OM membrane nano-discs were prepared as described above. Only the sucrose gradient used for membranes separation (IM and OM) was adjusted to the strain to 73% (w/v), 45% (w/v) and 20% (w/v) sucrose.

### NMR

NDs samples in Buffer B (100 mM NaCl, 25 mM Tris pH 7.4) were lyophilized and resuspended is 100% D_2_O. Around 3mg of sample was sedimented into a 1.3 mm ssNMR rotor at 68000 g for 16 h. Spectra acquisitions were setup using ssNMRlib suite^35^ for hard ^1^H and ^13^C pulses calibration as well as INEPT delays and cross-polarisation parameters, on a Bruker 600 MHz spectrometer equipped with MAS 1.3 mm HCN probes at 55 kHz rotation and 50°C. hCH CP and hCP INEPT were typically recorded in 24-36 hours. O157 O-antigen was assigned on ND-O157_pur_ sample using ^13^C (Single Quantum-J) and ^1^H detected experiments hCCH, hCH. Assignment of ^13^C O157 LPS solubilised in detergent micelles was performed on purified O157 LPS at 4mM in 60mM DHPC detergent in 100% D2O and buffer B at 50°C (^13^C-^1^H HSQC, HCCH TOCSY). All spectra were calibrated with respect to methanol (^1^H 2.225 ppm ^13^C 31.45 ppm), processed using TopSpin 3.5 software and analyzed using CcpNmr analysis 2.42 software.

### Bio-Layer Interferometry

ND-R1_pur_ and ND-R1_OM_, in PBS buffer pH 7 (euromedex), were biotinylated non-specifically on carboxylic groups using EDC and Biotin-LC-Hydrazide. From the approximate SMA concentration obtained from UV spectra at 260 nm, the EDC concentration was adjusted to react at maximum with 3% of SMA carboxylic groups. Typically for a ND sample with an OD_260nm_ of 0.2 about 15 µM EDC and 36µM Biotin-LC-Hydrazide were incubated for 2h at 25°C, then dialyzed against PBS.

The interferometry measurements were carried out on an octet Red96 (Fortébio) using streptavidin coated biosensors (SA). For interaction with MGL, the biotinylated nanodiscs were immobilized in HEPES buffered saline buffer, 0.02 % Tween20 to reach about 0.3 nm of response. The functionalized sensors were next equilibrated in interaction buffer (50 mM Phosphate 150 mM NaCl pH 8), then immersed in the wells containing MGL at concentrations ranging from 10 to 75 nM. MGL was biotinylated on carboxyls with EDC and Biotin Hydrazide. Protein at 2 mg/mL in buffer (150 mM NaCl 50 mM Phosphate pH 8) was treated with 1.25 mM Biotin LC-Hydrazide (Thermofisher) and 5 mM EDC for two hours then dialyzed against the initial buffer. Proteins were immobilized at 5µg/ml in HEPES buffered saline buffer, 0.02 % Tween20 to reach about 1 nm of response. The functionalized sensors were equilibrated in interaction buffer (50 mM phosphate 150 mM NaCl pH 8), then immersed in the wells containing R1 LOS vesicles at concentrations ranging from 0.7 to 15 µM. For interaction with polymyxins ND-R1_pur_ and ND-R1_OM_ were immobilized in PBS buffer to match conditions already described_32_. Immobilization level was 0.8 nm for all SA tips. All BLI data were processed and analyzed with Fortébio software after subtraction of the signal of a naked biosensor (SA) immersed in the same solution of analyte to remove the contribution of non-specific interactions. All Kd fittings were performed with a standard 1:1 model with simultaneous fitting of all curves shown (association and dissociation).

### AFM

AFM imaging was performed using a multimode 8 atomic force microscope with a Nanoscope V controller (Bruker). Imaging was performed in air using the PeakForce Tapping mode with a ScanAsyst Air cantilever (k = 0.4 N/m, Fq = 67 kHz). Images were recorded with a scan size of 2 × 2 μm^2^, at a line rate of 1 Hz and a resolution of 512 × 512 px^2^, using a semi-manual ScanAsyst control of the setpoint and/or gain. Data corrections include a plane leveling followed by a median line leveling using Gwyddion^36^. Stripe noise has been evaluated using DeStripe^37^ and visibility improvement has been tested using the Laplacian Weight Filter^38,39^.

All samples were loaded on a freshly cleaved mica surface coated with Ni^2+^ ion as follows. First, a 3 µl NiCl_2_ at 2mM was incubated for 2 min followed by drying under a gentle nitrogen flow. Then, 3 µL of a 30 to 120µg/ml of ND solution was applied and deposited for 2 min on the Ni-coated mica surface and dried under a nitrogen flow. Individual MGL proteins were diluted to 4.7 µg/ml prior to deposition on Ni-coated mica. The complex of MGL (4.7 µg/ml) and ND-R1pur (34 µg/ml) was incubated for 10 min before deposition.

Height and surface area were computed with Gwyddion using a standard grain analysis with a threshold segmentation. Individual grain heights were computed by subtracting the background height from the maximum height of the segmented grains. Grains were selected using a threshold of 1/3 of the maximum image height while the threshold for the background was the lower 10%. Grains were filtered by removing grains of smaller size (< 400 nm^2^ surface area), a value corresponding to the lower limit of the global average size of the smallest nanodiscs observed in this study. Height measurements were performed using Gwyddion’s “Distribution of various grain characteristics” tool, exported as text data, and further processed in Excel. Height and surface area were obtained from two different images.

### MGL-ECD expression & and purification

Human MGL isoform 2 extracellular domain (residues Q61-H292 Uniprot Q8IUN9-2) with an N-terminal Strep-tag II and a factor Xa cleavage site (MASWSHPQFEKIEGRGGG) was expressed and purified as already reported after refolding^21,23^.

### Dynamic light Scattering

DLS data of NDs were recorded at 25°C on Wyatt dynapro nanostar. For each sample 10 acquisitions of 5 sec were recorded and analyzed in percentage of mass.

### Quartz crystal microbalance with dissipation (QCM-D)

Quartz crystal microbalance with dissipation (QCM-D) experiments were carried out on a Q-Sense E4 instrument from Biolin Scientific (Gothenburg, Sweden). The quartz crystals were coated with 100 nm gold substrate (Model QSX 301) and fundamental resonance frequency of 4.95 MHz. Prior to use, the crystals were rinsed with a sequence of organic solvent (chloroform, ethanol, acetone) and water under sonication for 30 minutes, dried under nitrogen stream and treated by UV-ozone for 30 minutes. The baseline of QCM-D measurement was measured in the same buffer used for the experiments: 25 mM Tris, 100 mM NaCl, pH 7.4 for the study of MGL binding, and 50 mM Phosphate, 150 mM NaCl, pH 8 for the PmB adsorption studies. After establishing a stable baseline signal, a solution of ND-R1_pur_ (0.5 mg/mL) was injected at a flow rate of 100 µL/min. The protein was injected at a concentration of 12 µM in the same buffer as for the preparation of the lipid bilayer. The same holds for the injection of PmB, which was prepared at a concentration of 70 µM, in reference to previous studies on model bacterial membranes^40^. Data were collected using the Q-Soft software, provided by the company, and converted into data files with QTools.

## Supporting Information

Supplementary figures and tables are provided.

## Author Contributions

CL, SM, JLP designed the research; CL, SM, JLP and MA wrote the manuscript; CL, SM, JLP, MA, JMT, MT, MT, MM, FF performed the research; CL, SM, JLP, MA, JMT analyzed the data.

## Notes

The authors declare no competing financial interests.

## ACKNOWLEDGMENT

We thank the Agence Nationale de la Recherche (ANR) PIA for Glyco@Alps (ANR-15-IDEX-02) for the support of M.M., the Mizutani foundation for glycosciences (N° 230030) for their support of C.L. and the Grenoble Alliance for Integrated Structural and Cell Biology (ANR-17-EURE-0003) for funding M.A. This work used platforms of the Grenoble Instruct-ERIC center (ISBG; UAR 3518 CNRS-CEA-UGA-EMBL) within the Grenoble Partnership for Structural Biology (PSB), supported by FRISBI (ANR-10-INBS-0005-02). C.L. was granted access to the CCRT High-Performance Computing (HPC) facility under the Grant CCRT2023-lagurice awarded by the Fundamental Research Division (DRF) of CEA. This work acknowledges the AFM platform at the IBS. We thank B. Bersch for stimulating discussions and A. Vallet for technical help. This project has received funding from the European Research Council (ERC) under the European Union’s Horizon 2020 research and innovation program under grant agreement no 851356 to R.M.

