## Supplementary for "Lipopolysaccharides nanodiscs, a biomimetic platform to study bacterial surface"

### Supplementary information for : Lipopolysaccharides nanodiscs: a biomimetic platform to study bacterial surface

ND-O157pur

| | Radius (nm) | %Pd | Mw-R (kDa) | %Intensity | %Mass |
|---|---|---|---|---|---|
| Peak 1 | 11.933 | 27.9 | 1113 | 51.2 | 78.1 |
| Peak 2 | 122.758 | 13.8 | 260100 | 46.8 | 19.8 |

ND-O157OM

| | Radius (nm) | %Pd | Mw-R (kDa) | %Intensity | %Mass |
|---|---|---|---|---|---|
| Peak 1 | 8.8 | 13.0 | 550 | 13.9 | 83.5 |
| Peak 2 | 33.1 | 38.1 | 12140 | 81.5 | 12.4 |

| **AFM Surface Measurement** | | |
|---|---|---|
| | **ND-O157pur** | **ND-O157OM** |
| Number of values | 656 | 345 |
| Minimum (nm$^2$) | 400 | 404 |
| 25% percentile | 490 | 660 |
| **Median** | **626** | **934** |
| 75% percentile | 831 | 1415 |
| Maximum | 15000 | 43700 |
| **Mean** | **685** | **1124** |
| Standard deviation | 234 | 641 |

**Figure S1** *Left:* Dynamics light scattering data for NDs-O157. The symbol “Pd” stands for the size polydispersity. *Right:* statistics of AFM surface measurements of NDs-O157 deposited onto mica surface. Surface data are sorted in ascending order and the 25$^{th}$ percentile is the value that separates the lowest 25% of the data from the remaining 75%. Mean surface values of 685 nm$^2$ and 1124 nm$^2$ correspond to about 30 and 38 nm diameter respectively for a perfect disk.

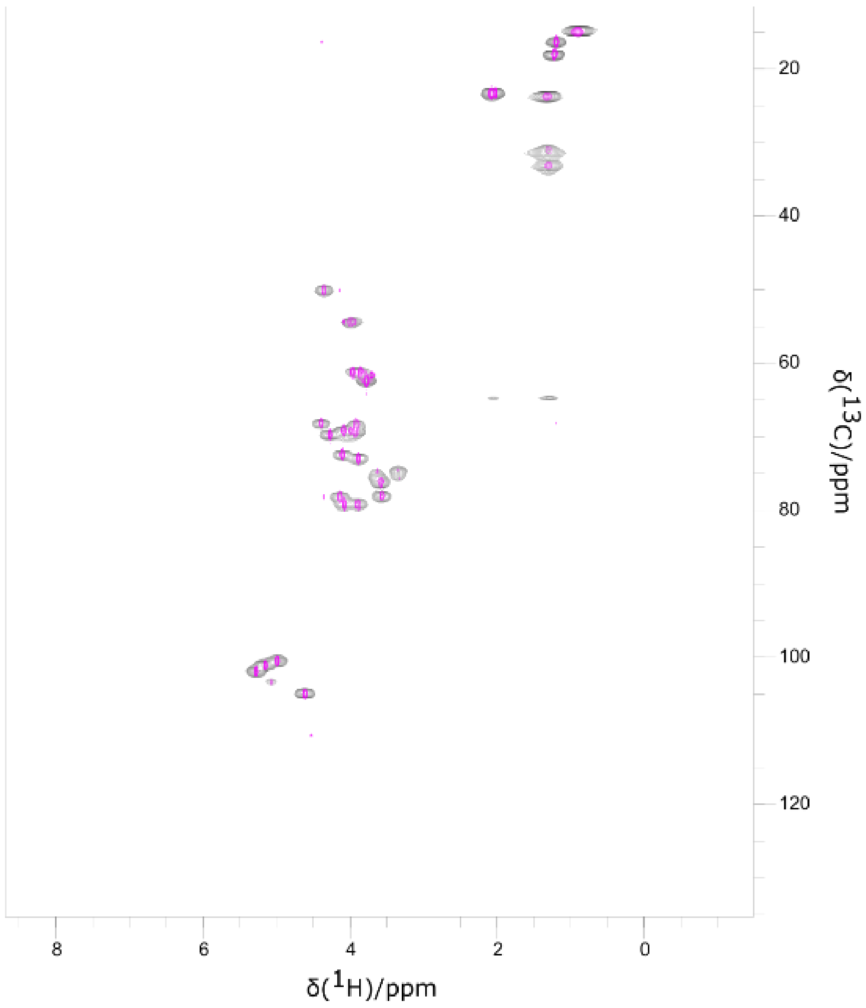


**Figure S2:** Solution NMR $^{1}H$-$^{13}C$ HSQC spectrum of ND-O157$_{pur}$(magenta) compared with its solid state $^{1}H$-$^{13}C$ INEPT at 55kHz (black) after sedimentation into the rotor. Both spectra were recorded at 50°C.

| | | C1 | C2 | C3 | C4 | C5 | C6 | C8 | H1 | H2 | H3 | H4 | H5 | H6 | H8 |
|---|---|---|---|---|---|---|---|---|---|---|---|---|---|---|---|
| W | GalN | 100,5 | 49,5 | 77,7 | 69,2 | 71,9 | 61,9 | 23,0 | 5,12 | 4,31 | 4,10 | 4,21 | 4,07 | 3,75 | 2,04 |
| X | Glc | 104,4 | 74,2 | 74,8 | 77,6 | 75,6 | 60,6 | | 4,58 | 3,31 | 3,59 | 3,55 | 3,55 | 3,86 | |
| Y | Fuc | 99,9 | 67,8 | 72,4 | 78,7 | 67,6 | 15,9 | | 4,96 | 3,86 | 3,86 | 3,87 | 4,36 | 1,17 | |
| Z | PerN | 101,3 | 78,6 | 68,6 | 53,8 | 68,8 | 17,5 | | 5,24 | 4,06 | 3,90 | 3,96 | 4,07 | 1,20 | |
| Z' | PerNTer | 102,7 | | | | | | | 5,01 | 4,02 | | | | | |

**Figure S3:** Assignment of O157 O-antigen in ND-O157$_{pur}$ by ssNMR. Top: $^{13}C$-$^{13}C$ Single Quantum J-correlation experiment of ND-O157$_{pur}$ nanodiscs at 15kHz, 50°C. Bottom: $^{1}H$-$^{13}C$ INEPT and hCCH correlation spectra of ND-O157$_{pur}$ at 55kHz by proton detection.

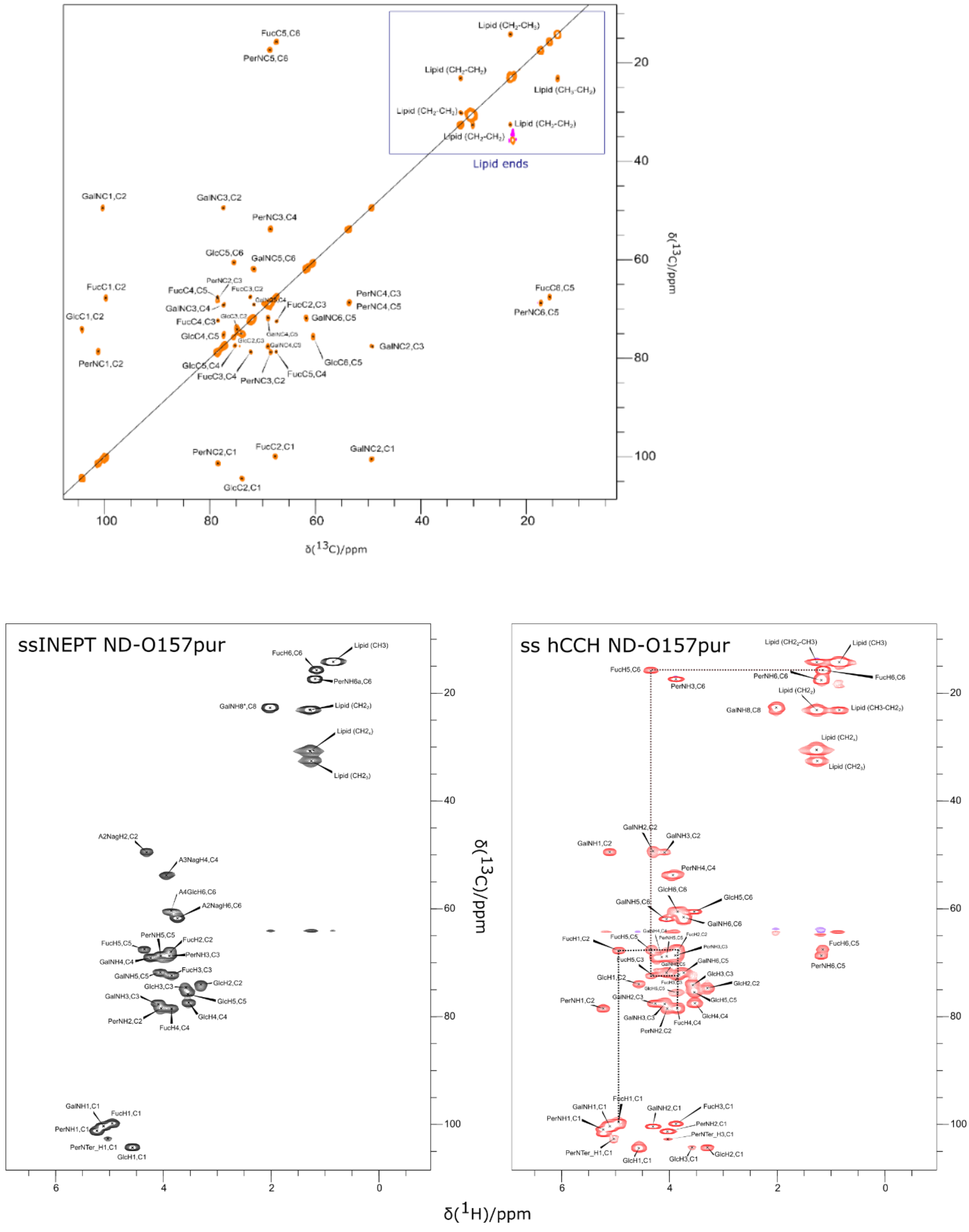

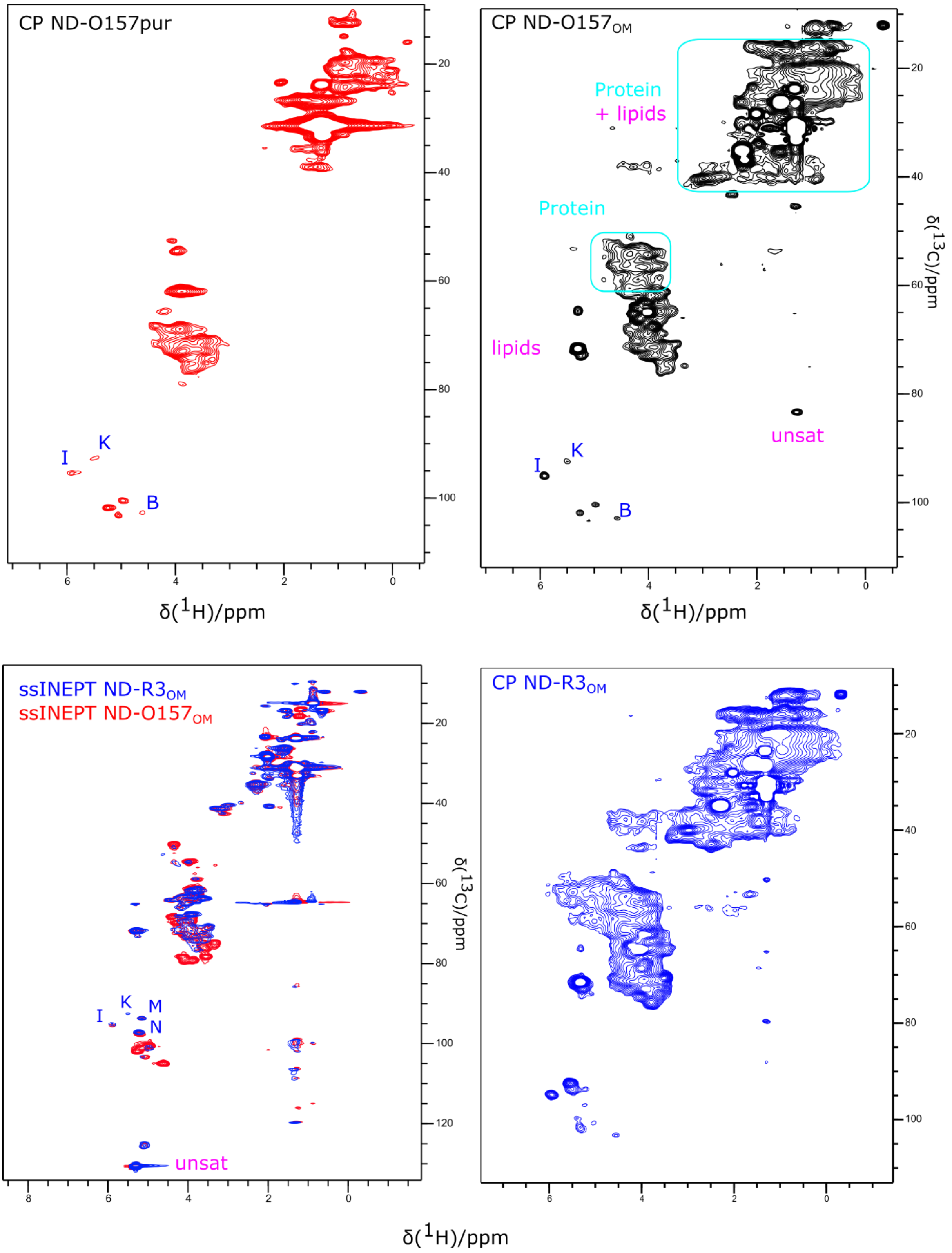


**Figure S4**: Supporting solid state NMR spectra of ND-O157pur, ND-O157$_{OM}$ and ND-R3$_{OM}$ by $^{1}$H detection at 55 kHz and 50 °C.

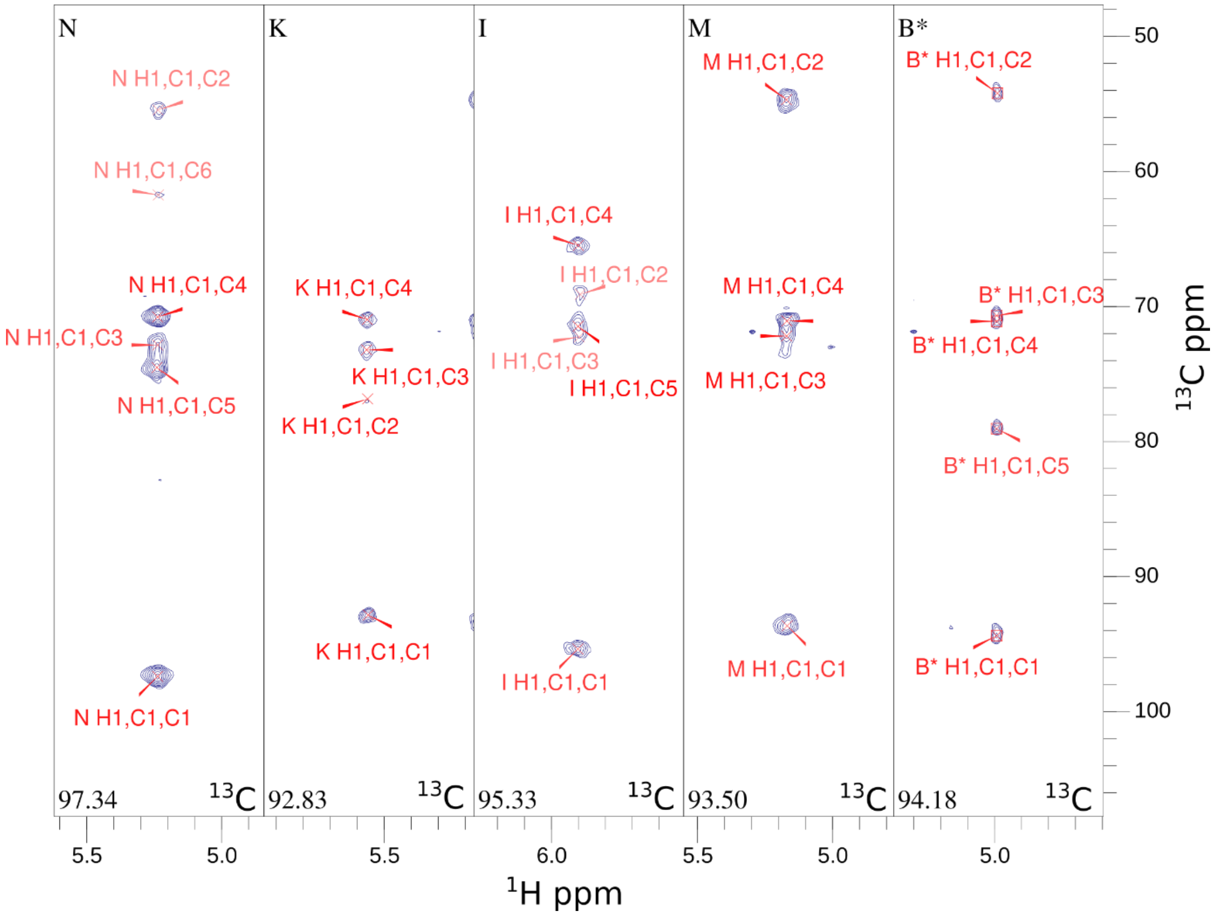


| | | C1 | C2 | C3 | C4 | C5 | C6 | H1 | H2 | H3 | H4 | H5 | H6 |
|---|---|---|---|---|---|---|---|---|---|---|---|---|---|
| N | GlcN | 97,3 | 55,4 | 72,9 | 70,7 | 74,5 | 61,8 | 5,23 | 3,34 | 3,57 | 3,47 | 3,78 | |
| K | Glc | 92,8 | 76,8 | 73,1 | 71,0 | 76,2 | 62,1 | 5,56 | 3,76 | 3,94 | 3,52 | | |
| I | Gal | 95,3 | 69,2 | 72,2 | 65,5 | 71,3 | 62,3 | 5,9 | 3,8 | 4,25 | 4,28 | 4,3 | 3,79,3,83 |
| M | GlcNAc | 93,5 | 54,7 | 72,3 | 71,3 | 73,3 | 62,0 | 5,16 | 4,04 | 3,86 | 3,59 | | |
| B* | GlcN | 102,9 | 52,6 | 75,7 | 78,2 | 81,2 | 69,0 | 4,61 | 4,12 | | | 3,93 | 4,19 |

**Figure S5.** Assignment of R3 Core Oligosaccharide from O157 LPS in detergent micelles by solution NMR. Strips of HCCH Tocsy ($^{13}C$-$^{1}H$) at anomeric groups of O157 LPS in DHPC detergent micelles. Table of the assigned sugars from Lipid A and core of R3.

A R1 LOS *E. coli F470 (R1)*

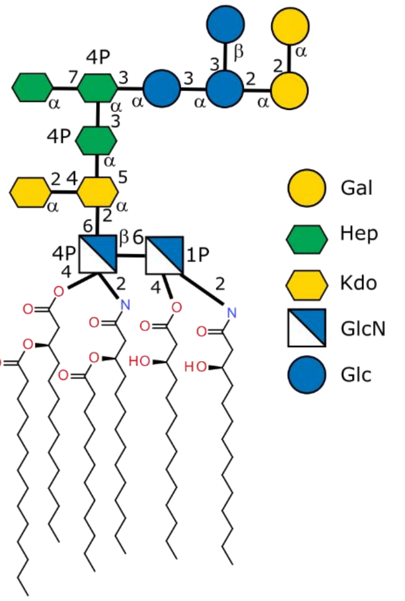


B

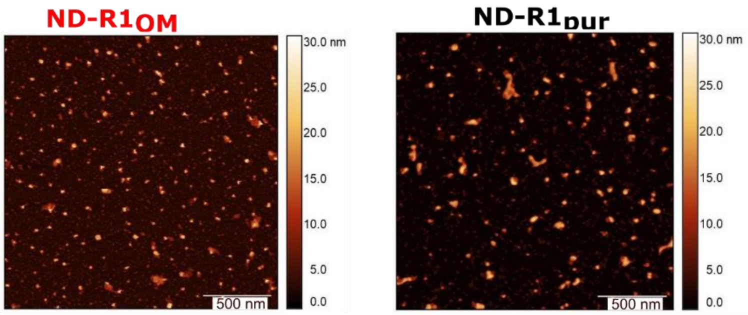


| AFM Surface Measurement | | |
|---|---|---|
| | **ND-R1pur** | **ND-R1OM** |
| Number of values | 278 | 360 |
| Minimum (nm2) | 416 | 406 |
| 25% percentile | 783 | 614 |
| **Median** | **1340** | **855** |
| 75% percentile | 2153 | 1255 |
| Maximum | 14000 | 4950 |
| **Mean** | **1712** | **1028** |
| Standard deviation | 1396 | 604 |

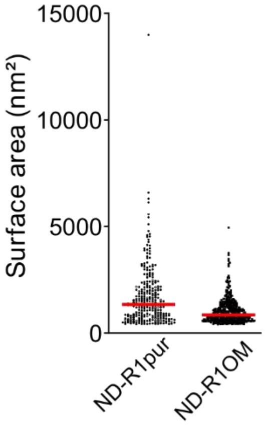


C ND-R1pur

| | Radius (nm) | %Pd | Mw-R (kDa) | %Intensity | %Mass |
|---|---|---|---|---|---|
| Peak 1 | 9.1 | 32.8 | 594 | 93.4 | 98.6 |
| Peak 2 | 221.5 | 6.5 | 1034850 | 2.3 | 0.5 |

ND-R1OM

| | Radius (nm) | %Pd | Mw-R (kDa) | %Intensity | %Mass |
|---|---|---|---|---|---|
| Peak 1 | 6.5 | 20.5 | 264 | 14.8 | 85.3 |
| Peak 2 | 20.9 | 34.4 | 4118 | 85.2 | 14.7 |

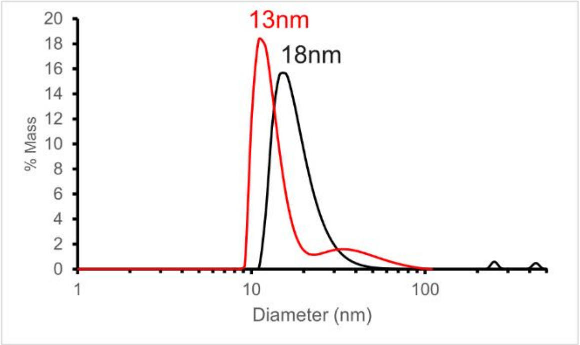


**Figure S6.** Characterization of ND-R1$_{pur}$ and ND-R1$_{OM}$

The structure of LOSR1 is shown in A, with the AFM imaging, surface measurement statistics and distribution in B and DLS statistics and distribution in percentage of mass in C with ND-R1$_{OM}$ in red.

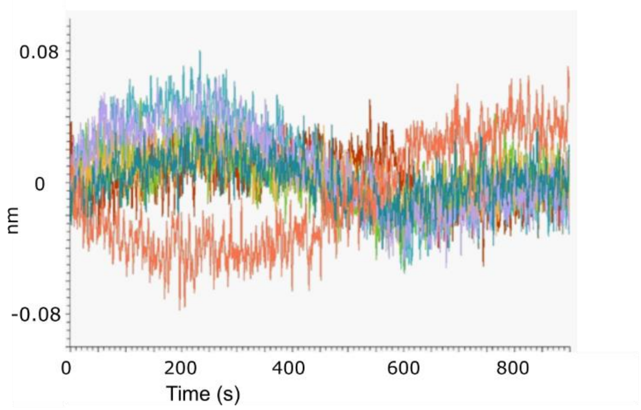


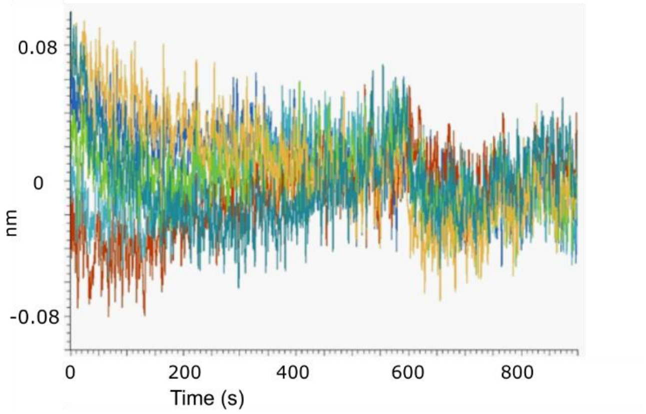


| | $K_D$ (M) | $K_D$ Error | $k_a$ (1/Ms) | $k_a$ Error | $k_{dis}$ (1/s) | $k_{dis}$ Error |
|---|---|---|---|---|---|---|
| **MGL ND-R1$_{purBiot}$** | 3,75 E-09 | 7,47 E-11 | 1,72 E+04 | 5,76 E+01 | 6,45 E-05 | 1,27 E-06 |
| **MGL ND-R1$_{OMBiot}$** | 3,18 E-09 | 4,57 E-11 | 2,44 E+04 | 6,04 E+01 | 7,75 E-05 | 1,10 E-06 |

**Figure S7. BLI of MGL interaction with ND6R1purbiot and ND-R1OMBiot.**

Residuals of fitting from figures 3A,B for ND-R1$_{pur}$(top) and ND-R1$_{OM}$ (middle) interaction with the fitted kinetic values below.

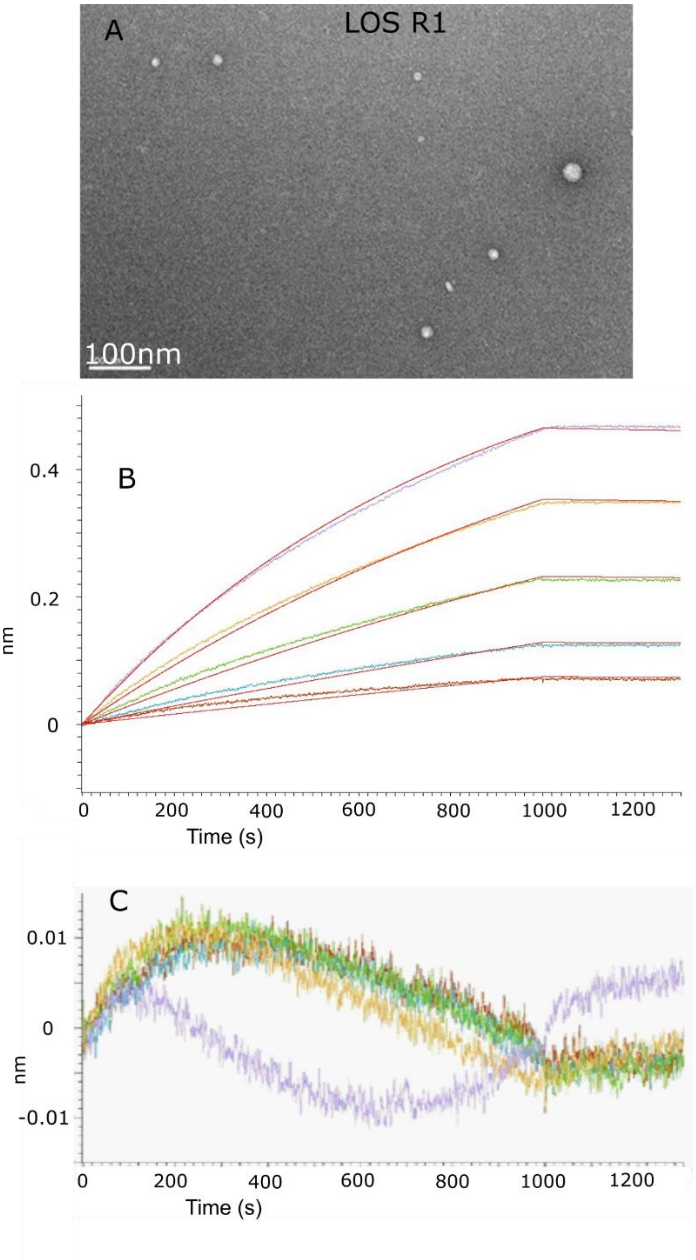


**Figure S8**. A, Negative stain electron micrograph of R1 LOS vesicles used for interaction with immobilized MGL (top). B, Increasing R1 LOS vesicle concentrations were flowed (0.7, 1.2 1.9, 3.2, 5.4 µM) over Immobilized MGL and monitored by BLI (left). The apparent $K_d$ fitted from simultaneous fitting of association/dissociation of these sensorgrams was 129 nM +/-3. C, The residual between fit and raw data is shown with fitting results below.

| | $K_d$ (M) | $K_d$ Error | $k_a$ (1/Ms) | $k_a$ Error | $k_{dis}$ (1/s) | $k_{dis}$ Error |
|---|---|---|---|---|---|---|
| **LOS-R1 MGLbiot** | 1,29 E-07 | 3,23 E-09 | 2,20 E+02 | 5,43 E-01 | 2,83 E-05 | 7,06 E-07 |

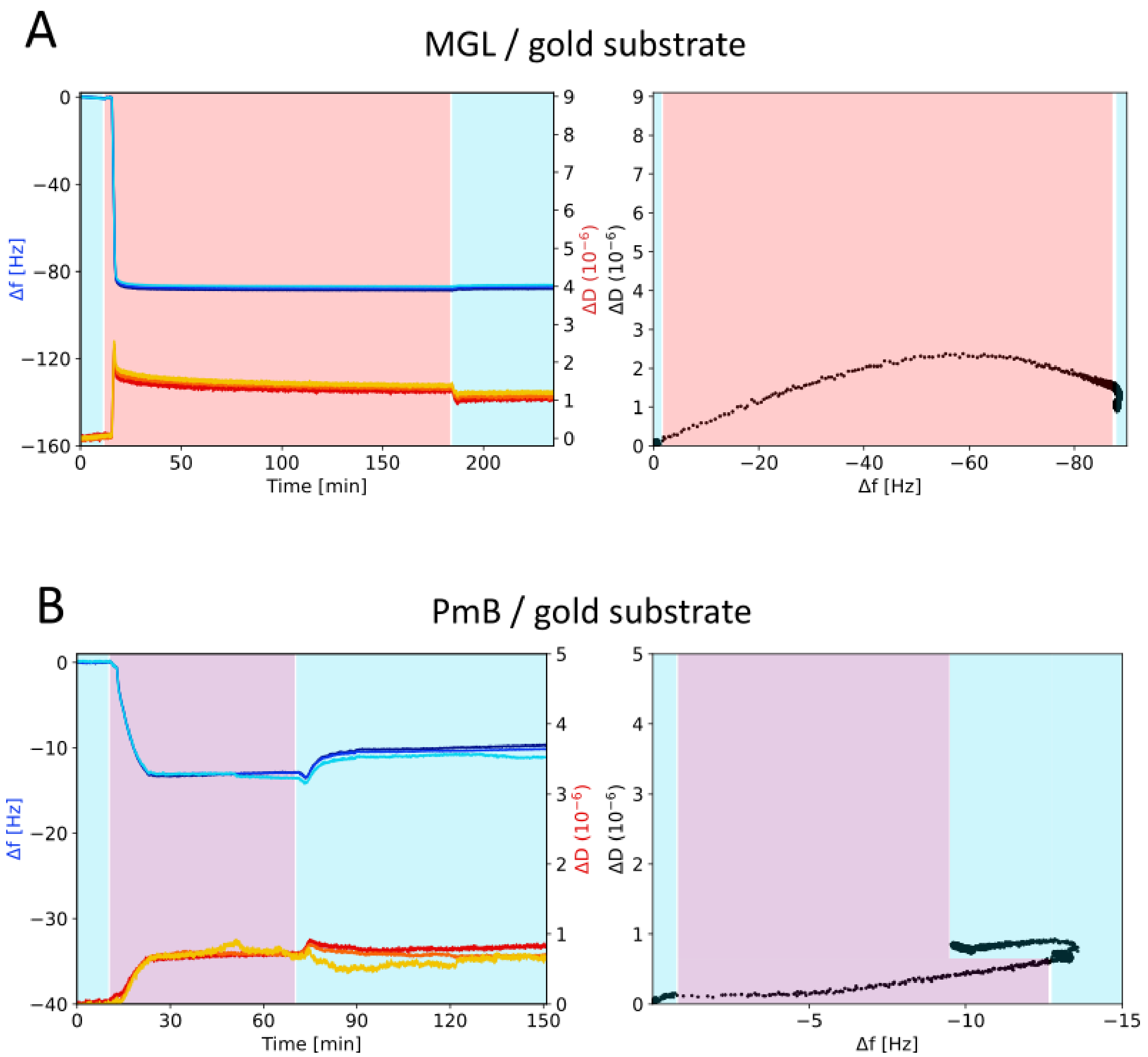


**Figure S9**. QCM-D: Adsorption studies of MGL (A) and PmB (B) onto bare gold substrate.

*Left*: Δf and ΔD signals registered during the adsorption of the molecules (A: MGL; B: PmB) onto the gold substrate. *Right*: ΔD vs. Δf plots to directly compare the variation of the viscoelastic properties (film softness/rigidity) with the variation of the adsorbed mass.

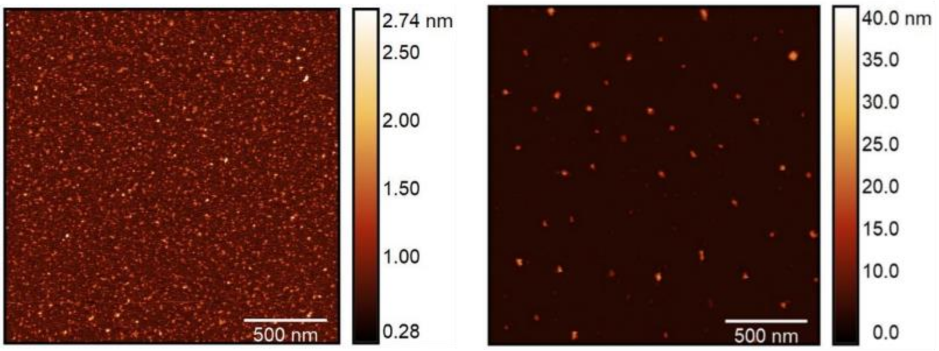


| Height (nm) | **ND-R1pur (above right)** | **+MGL (Fig. 3C)** |
|---|---|---|
| Number of values | 108 | 163 |
| Minimum | 8.4 | 10.8 |
| Maximum | **24.1** | **32.7** |
| **Mean** | **16.2** | **20.4** |
| Standard deviation | 3.5 | 4.8 |

**Figure S10**: Complement of Fig. 3C with AFM imaging of MGL alone (left) and ND-R1$_{pur}$ alone (right) with the statistics of AFM height measurements with/without MGL addition.
